# Auditory Brainstem Response as an Objective Measure of Temporal Resolution

**DOI:** 10.1101/2022.10.23.513412

**Authors:** Varsha M Athreya, Animesh Barman

## Abstract

**Background and Aims:** Temporal resolution is assessed using behavioral tests, which are highly affected by extraneous variables. We explored the relationship between behavioral Across-Channel Gap Detection Threshold (AC GDT) and different temporal parameters of an objective measure, Auditory Brainstem Response (ABR), to account for the extraneous variables.

**Settings and Design:** We conducted an experimental study on thirty native Kannada-speaking adults with normal hearing sensitivity in the age range of 18-25 years.

**Materials and Methods:** We estimated the Gap detection threshold (GDT) using an across-channel (AC) paradigm, and Auditory Brainstem Response (ABR) was recorded at 80 dBnHL for three repetition rates of 11.1, 30.1, and 90.1/sec.

**Statistical Analyses used:** Normality testing, the Friedman test, post-hoc analysis by Wilcoxon Signed-Rank test, along with descriptive statistics was performed using SPSS v20 (IBM Statistical Package for Social Sciences, New York, USA).

**Results:** The results showed a significant positive correlation between AC GDT scores and latency of wave I for the repetition rates of 11.1 and 30.1/sec and latency of wave V at 30.1 and 90.1/sec. There was a negative correlation (but not significant) between AC GDT scores and the slope of wave V across the repetition rates.

**Conclusions:** The results suggest a relationship between the behavioral and electrophysiological measures of temporal processing. Measuring the latency of wave I and wave V of ABR would give an estimate of their AC GDT scores, especially in difficult-to-test populations.

**Key messages:** To test temporal resolution abilities in individuals with normal hearing sensitivity, we can use an objective measure of latency of wave I and wave V of ABR. ABR can be highly useful in reducing testing time and obtaining reliable estimates in children and individuals with associated disorders like autism, below-average intelligence, and so on.

## Introduction

Communication in daily life is beyond detecting sounds and involves using different processes to extract the key features of the auditory input. One of these processes is temporal processing, which gives information regarding the coding of time-related changes in the auditory input and is highly important in the perception of speech in noise.^[1]^ The utilization of temporal processing for speech perception is highly variable across individuals.^[2]^ A commonly used behavioral test to assess temporal resolution is the Gap Detection Test (GDT). Here, we used the across-channel GDT (AC GDT) paradigm, comparing time-related changes in neural activity across different auditory channels.^[3]^ Studies have also revealed that AC GDT correlates better with speech perception in noise.^[4]^ However, AC GDT is a behavioral measure affected by extraneous variables like attention, interest, procedural differences, etc.

To assess variation in temporal processing using an objective test, we utilized Auditory Brainstem Response (ABR). ABR requires high temporal precision, and the responses are a direct measure of the neural integrity of the anatomical region.^[5]^ We used a validated behavioral measure of AC GDT to identify the parameters of ABR as an objective measure of temporal resolution. The relationship between AC GDT values and temporal parameters of ABR obtained at different repetition rates was explored in the present study.

## Subjects and Methods

### Participants

We considered thirty native Kannada-speaking adults aged 18-25 years (M= 20.77 years) for the study. The inclusion criteria were bilateral normal pure tone thresholds (≤15 dBHL between 250 Hz to 8000 Hz), bilateral normal ‘A’ type tympanogram with reflexes present, and normal bilateral transient evoked otoacoustic emissions (TEOAE). Through a structured interview, we ruled out the possibility of ototoxicity and noise exposure. The study adhered to the Ethical guidelines for bio-behavioral research involving human subjects at All India Institute of Speech and Hearing, Mysuru.^[6]^All the tests were carried out in an air-conditioned sound-treated double room. The ambient noise level was within the permissible limits.^[7]^

### Procedure

#### Gap Detection Test

Gap Detection Threshold (GDT) was estimated using an across-channel paradigm. The stimuli used were narrow-band noise (NBN) centered at 1000 Hz (lead marker) and 2000 Hz (lag marker). The signal processing and stimulus presentation was generated instantaneously in Dell Inspiron 15 (Dell Technologies, Round Rock, USA) laptop loaded with Psycon v2.18 experimental software with a sampling rate of 44100 Hz, using AUX scripting.^[8]^ The lead marker’s rise time and the lag marker’s fall time were set to 30 ms. The duration of the lead marker was kept constant at 300 ms, with the duration of the lag marker varying from 230 ms to 300 ms to minimize cues toward the identification of the gap. The overall stimulus duration remained constant within and across the test procedure. The standard or the reference stimulus had a constant gap duration of one ms, whereas the test stimulus had a variable gap duration. To reduce the spectral splatter introduced by the abrupt onset and offset of the NBN, ramping of one ms was used before and after the gap^[1]^ *(Fig 1).* The test began with a gap size of 70 ms. In case of an incorrect response, the gap size was increased till the participant could identify the gap correctly in at least two consecutive trials. The gap size increased with every incorrect response and decreased with two correct responses, with the step sizes being seven and two ms above and near the threshold, respectively.

**Fig 1.**
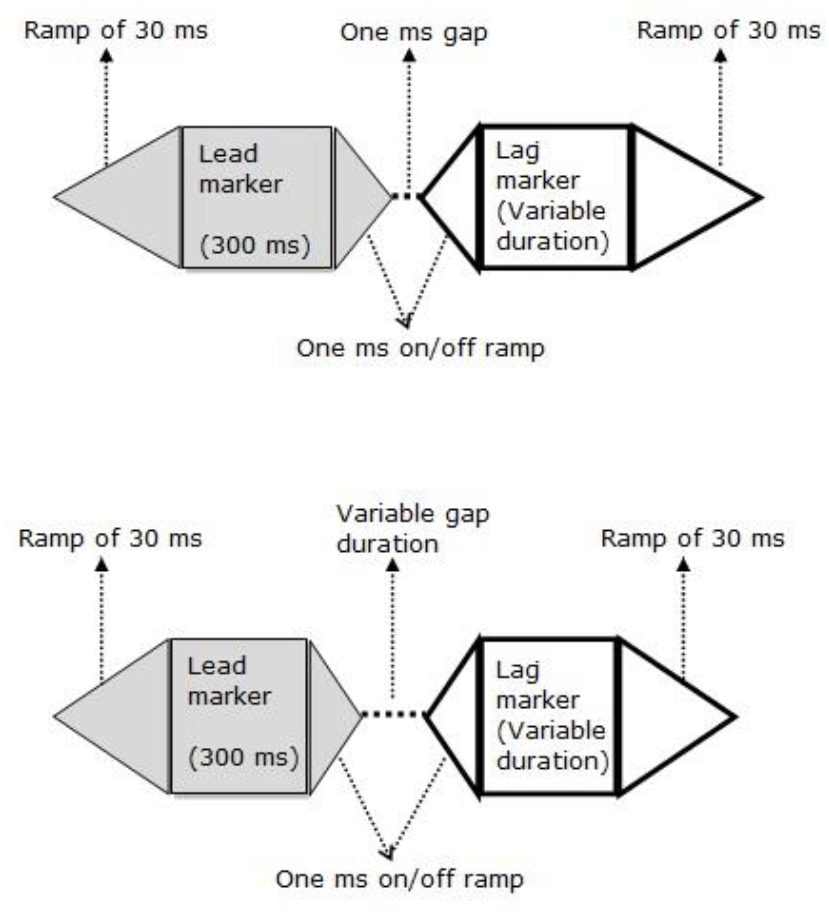
Schematic representation of the stimulus used for Across-Channel Gap Detection Testing.

**Fig 2.**
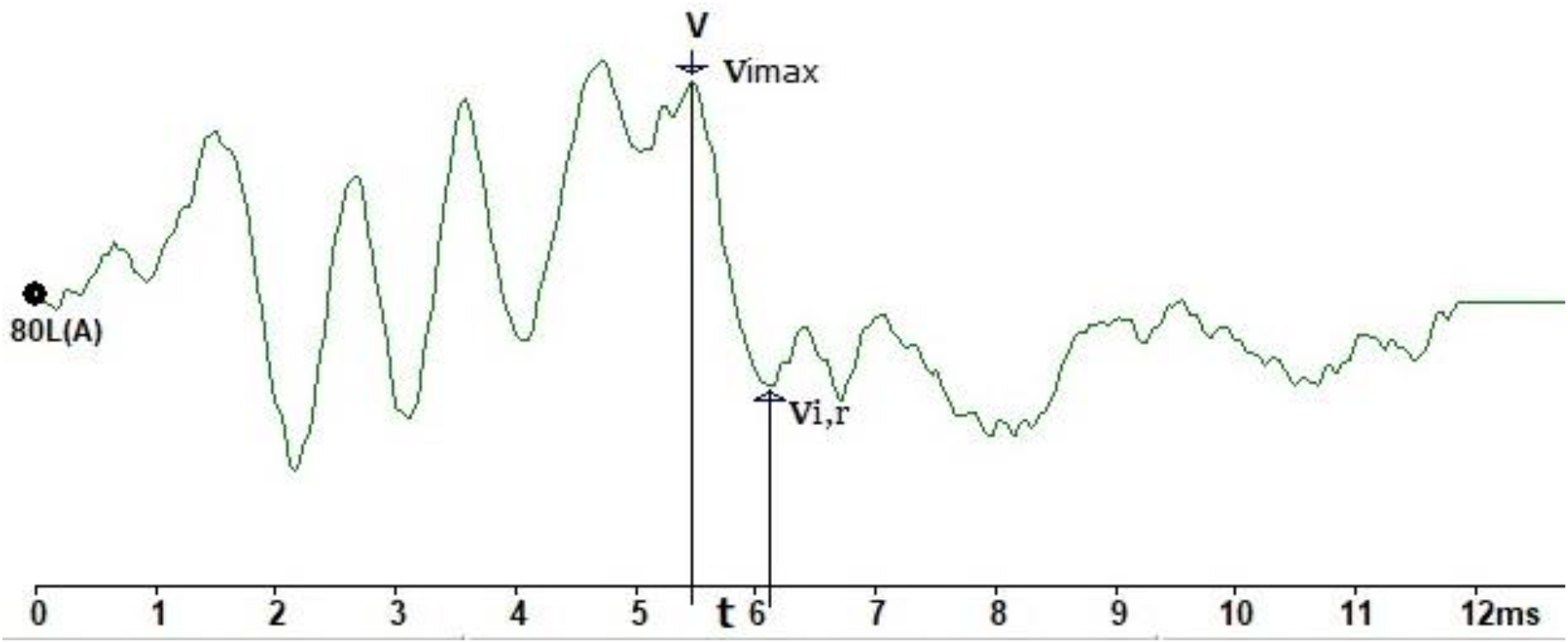
Representation of the calculation of slope of wave V of Auditory Brainstem Response.

We obtained the GDTs of both ears for all the participants. The stimulus was presented at the most comfortable level (MCL). Two down/ one up (2D1U) adaptive procedure with a hit rate of 70.7% was used to estimate the thresholds.^[9]^ The test was conducted using a three-intervals three-alternative forced-choice (3 AFC) method with an intra-block interval of 500 ms and inter-block interval of 1000 ms referenced to the subject’s response.

The test was terminated after eight reversals. The mean gap size of the last six intervals was calculated to obtain the Gap Detection Threshold. The above procedure was administered thrice for each ear, for every individual. The best GDT value amongst the last two trials was considered the AC GDT.

#### Auditory Brainstem Response (ABR)

ABRs were recorded using 100 μs rarefaction clicks presented through ER 3A (Etymotic Research, Inc., Elk Grove Village, USA) insert earphones at three repetition rates of 11.1, 30.1, and 90.1/sec. The stimuli were presented at 80 dB nHL. ABRs were elicited using IHS SmartEP windows, USB v4.0 (Intelligent Hearing Systems, Miami, USA). The recording parameters for a time window of 15 ms were bandpass filtering from 100 to 3000 Hz and a gain of 100000 times. The recordings were obtained with the non-inverting electrode placed at the vertex, inverting electrode on the mastoid of the test ear, and the ground electrode on the mastoid of the non-test ear. Two recordings of 2000 sweeps were obtained at each repetition rate, and their averaged waveform was used for analysis. ABRs were elicited for both ears separately for every participant.

#### Waveform Analysis

The waveforms were analyzed by two audiologists and the experimenter. The waves were marked where two out of the three examiners agreed upon. For the repetition rates of 11.1, 30.1, and 90.1/sec, (1) the latency of wave I and wave V and (2) the slope of wave V were analyzed. The slope of wave V was calculated using the decay of wave described by Gopal and Kowalski (1999)^[10]:^

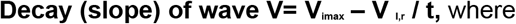

V_imax=_ Max peak potential of the wave
V_I,r_= Potential at the immediate succeeding right minima
t= Duration of the wave’s descent

#### Statistical Analysis

The statistical procedures were conducted using SPSS v20 (IBM Statistical Package for Social Sciences, New York, USA). Shapiro-Wilks test indicated non-normal distribution, and hence non-parametric tests - Friedman test, post-hoc analysis by Wilcoxon Signed-Rank test was used.

## Results

### AC GDT values

Descriptive analysis of AC GDT scores showed that the mean GDT value was 35.89 ms (SD: 11.98 ms) and the median was 33.50 ms. The thresholds ranged from 19 ms to 66.80 ms.

### Temporal parameters of ABR

Response rate measurement for the presence of wave I and wave V revealed a low rate of response of wave I, especially at higher repetition rates, whereas wave V was present in all individuals across the repetition rates *(Table 1).*

**Table 1.**
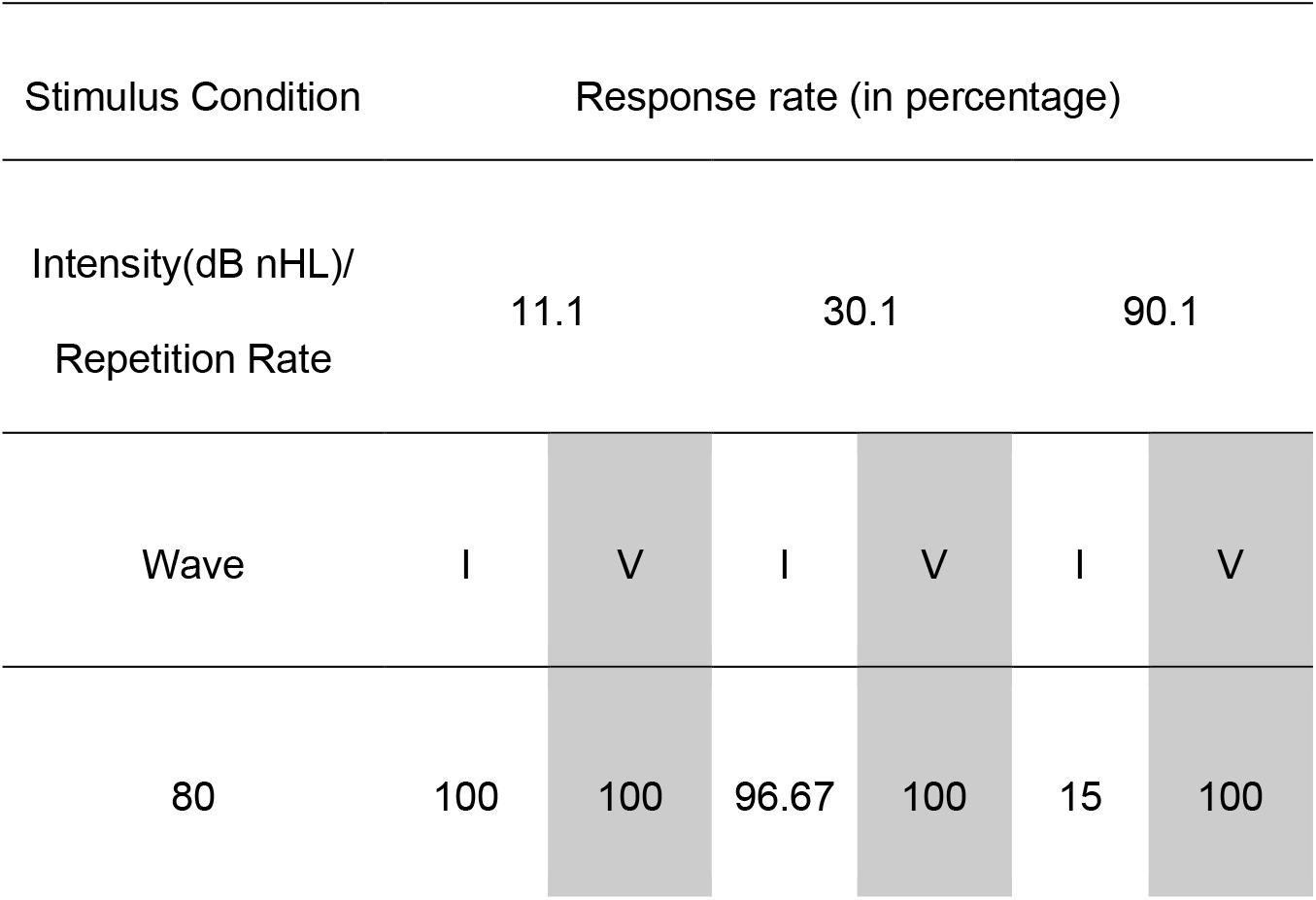
The response rate of wave I and wave V (in percentage) across the repetition rates of 11.1/sec, 30.1 /sec, and 90.1 /sec at 80 dB nHL

### Latency of wave I and wave V

Wave I showed an increase in latency with the increase in repetition rate. There was a greater increase in latency when the repetition rate changed from 30.1 to 90.1/sec *(Fig 3).* Friedman test revealed that the latency of wave I increased significantly with the increase in repetition rate at 80 dB nHL (χ^2^ (3) = 15.77, *p* = 0.00) with significant difference (*p*<0.01) amongst all the pairs of repetition rates as shown by Wilcoxon signed-rank post hoc test analysis. A significant difference was seen between 11.1 and 30.1/sec for wave I.

**Fig 3.**
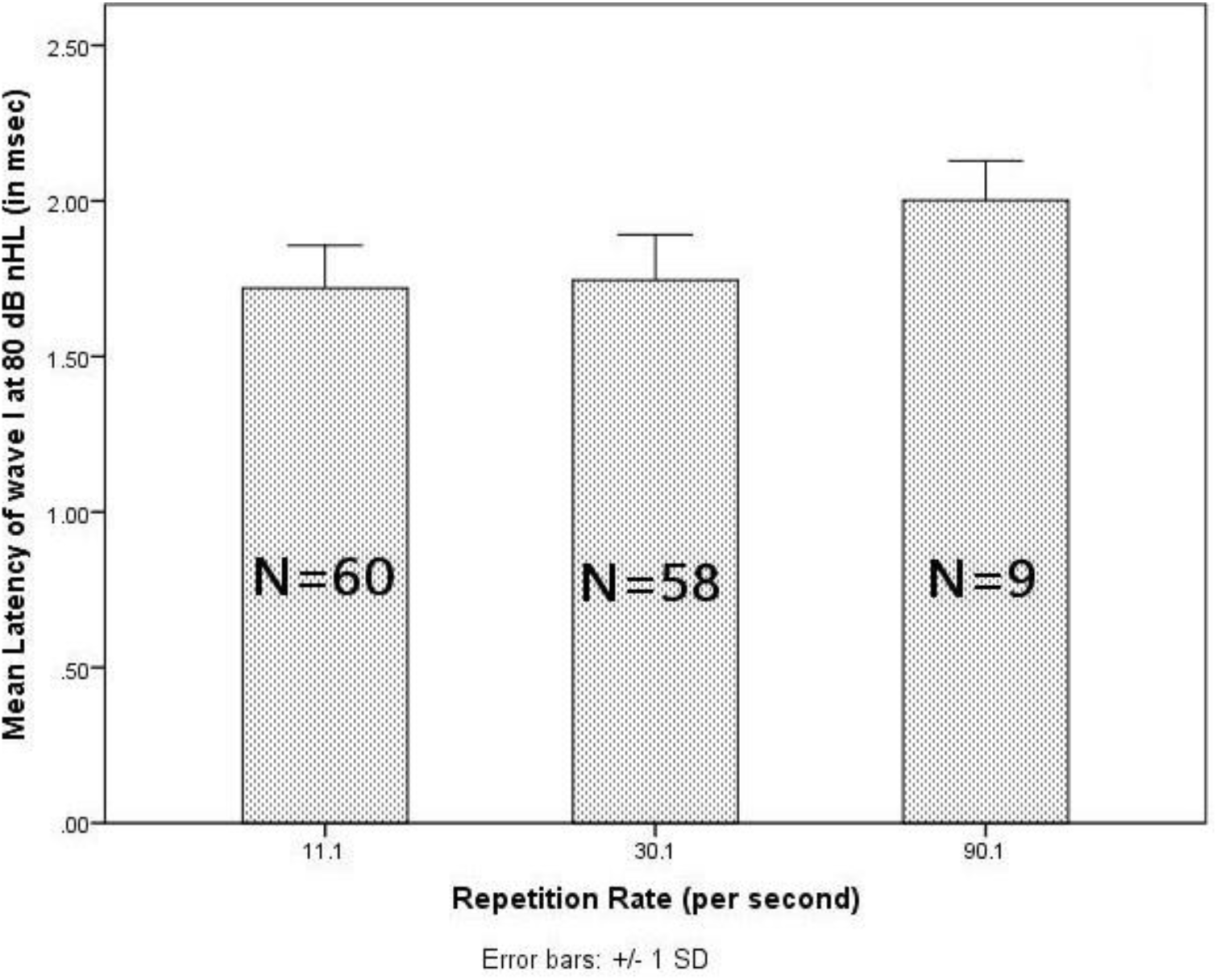
Latency of wave I of ABR obtained at 80 dB nHL across repetition rates

The latency of wave V increased with the increase in repetition rate (*Fig 4*). Friedman test showed that this increase in latency of wave V was significant across repetition rates, χ^2^(3) = 112.00, *p* = 0.00. Wilcoxon signed-rank test exhibited a significant difference (*p*<0.01) in the latency of wave V across all the pairs of repetition rates.

**Fig 4.**
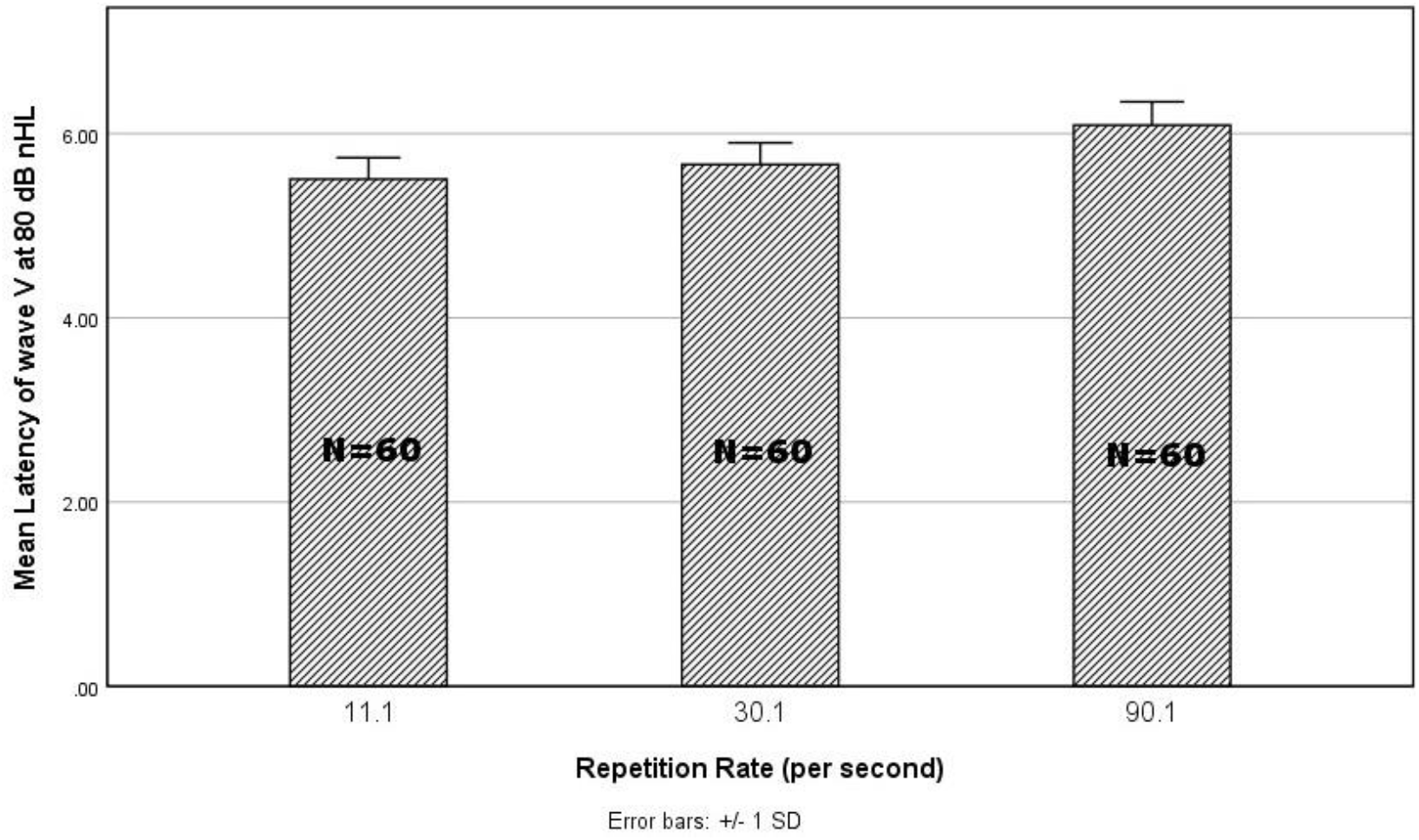
Latency of wave V of ABR obtained across repetition rates

### Slope of wave V

The mean and median slope of wave V decreased with the increase in repetition rate. A sharp wave V resulted in a higher slope value and a lower value for broader wave V. This temporal measure also exhibited large variations in individuals with normal hearing sensitivity (*Fig 5*). Friedman test revealed a significant effect of the slope of wave V across repetition rates, χ^2^(3) = 25.59, *p* = 0.00. Wilcoxon signed-rank test showed a significant difference (*p*<0.01) between all the repetition rates.

**Fig 5.**
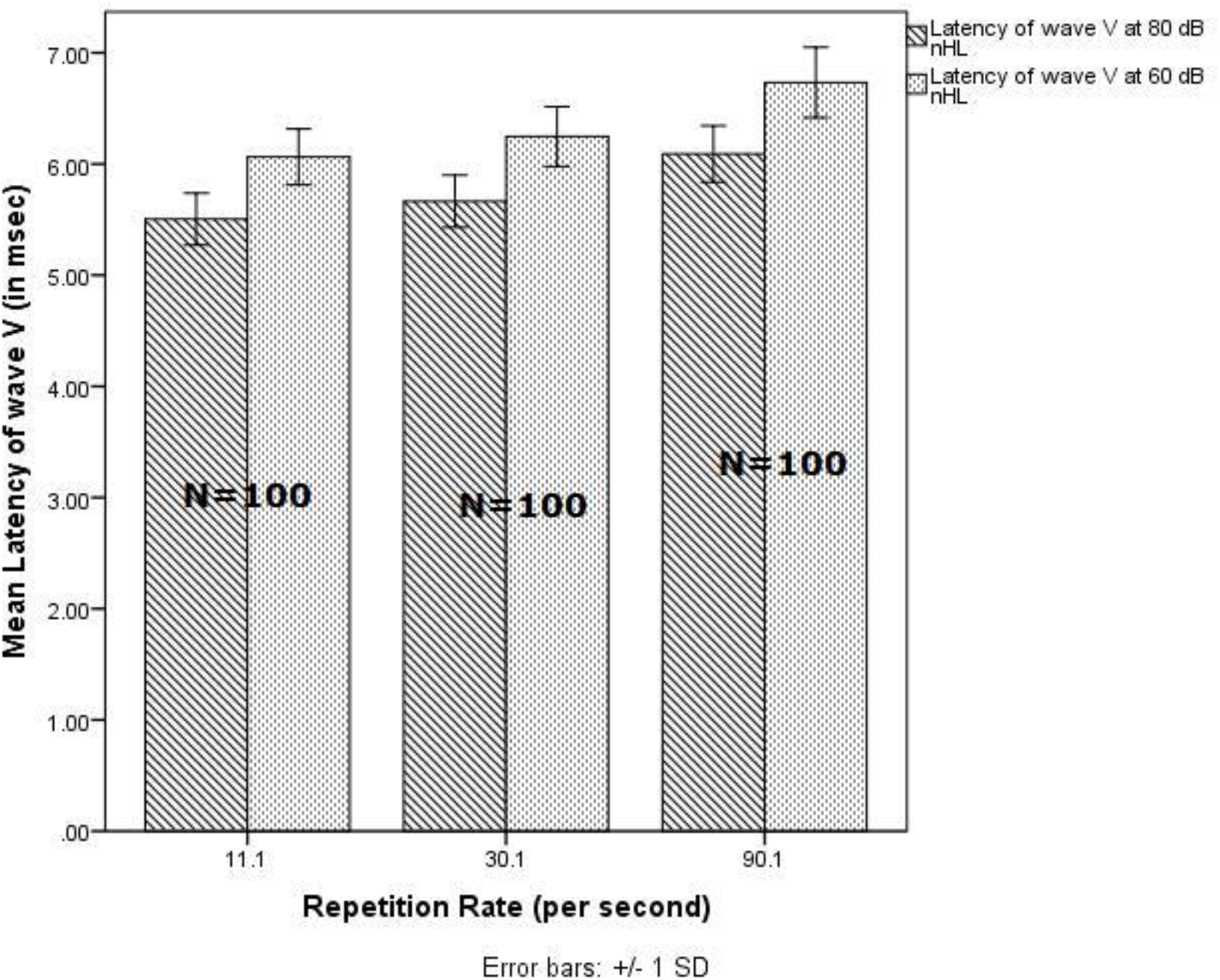
Slope of wave V of ABR obtained across repetition rates

### Correlational Analysis

The primary aim of the present study was to explore the relationship between the behavioral measure of temporal processing and its neural correlates at the brainstem level. Correlational analysis was performed using Spearman’s rank-order correlation between AC GDT values and the temporal parameters of ABR. A significant positive correlation was seen between AC GDT scores and the latency of wave I at 11.1 and 30.1/sec repetition rates and the latency of wave V for the repetition rates of 30.1 and 90.1/sec.

A negative non-significant correlation (*p* >0.05) was observed between AC GDT scores and the slope of wave V across the repetition rates *(Table 2).*

**Table 2.**
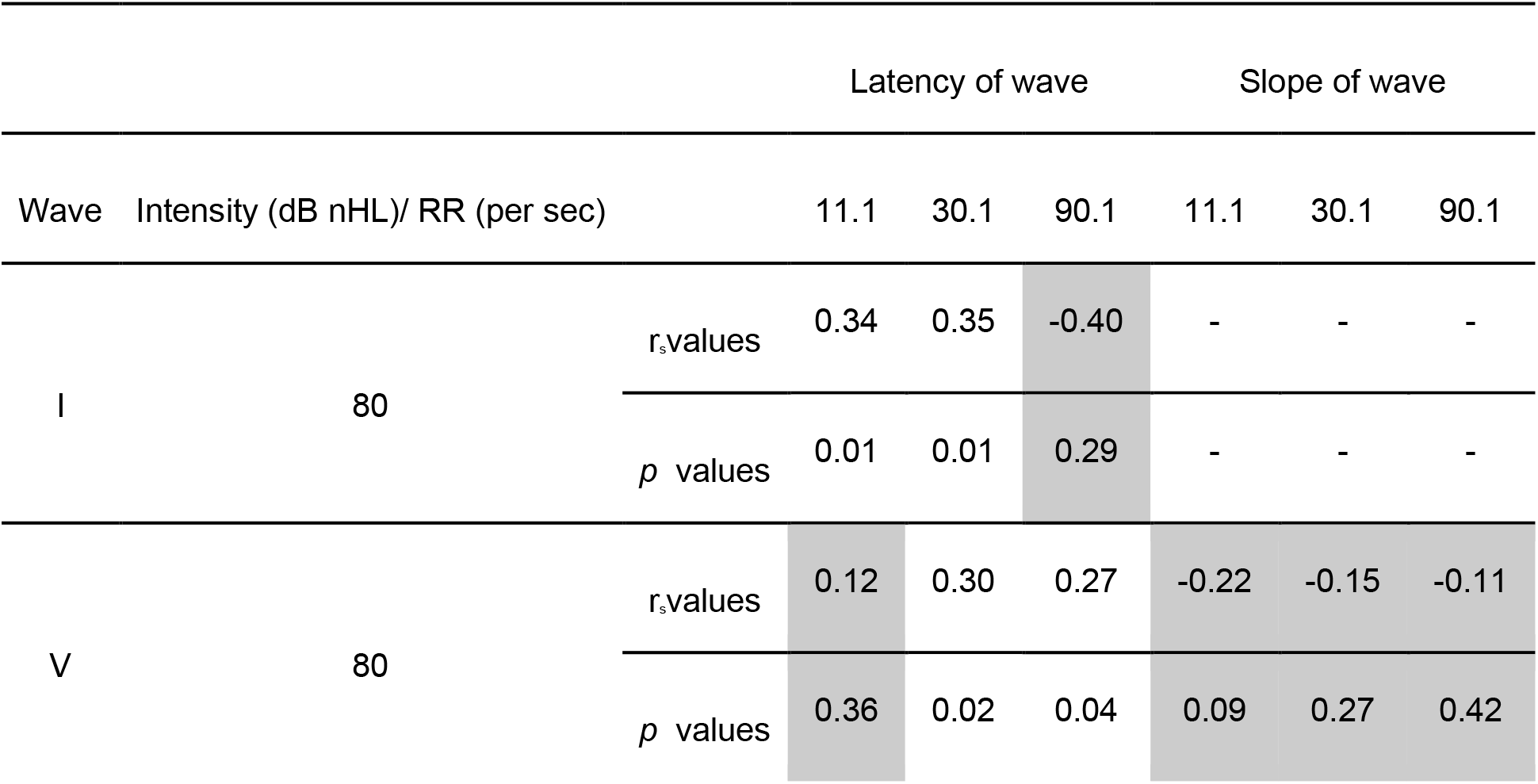
Results of Spearman’s rank-order correlation, r_S_, and significance level for AC GDT scores and latency of wave I and wave V, and slope of wave V across different repetition rates (dark-shaded boxes indicate no significant correlation).

## Discussion

### Variation seen in AC GDT scores

The data in the present study shows a large range of AC gap detection thresholds in agreement with previous studies.^[1, 11]^ AC GDT requires the activation of two perceptual neural channels. The offset and onset of the lead and lag marker change the deactivation and activation of neural activity across different auditory channels. Comparing timing between these two neural channels to detect the gap requires highly precise temporal resolution skills.^[1]^ There is a large variability in the use of temporal resolution skills in individuals with normal hearing sensitivity, which is seen in the wide range of AC GDTs.^[2]^

#### Latency of wave I and wave V

The results of the current study showed an increase in the latency of wave I and wave V with the increase in repetition rate, as reported previously.^[12, 13]^ With the increase in repetition rate, the neurons along the auditory pathway undergo adaptation, leading to an increase in latency of wave I (at the level of inner hair cells and auditory nerve junction) and wave V (up to inferior colliculus).^[12, 13]^ The firing rate of the individual neurons reduces with the prolonged presentation of stimulus or with a highly taxing stimulus to the auditory system, leading to increased wave I and wave V latencies.^[14]^

#### Slope of wave V

We observed that the slope of wave V decreased with the increase in the repetition rate. A higher value of the slope is indicative of a sharper wave V, whereas a broad wave V would have a lower value of the slope. A broad wave V can be attributed to wave V’s prolonged onset and offset.^[13]^ With an increased repetition rate, the adaptation of auditory neurons leads to reduced neural activity impacting the wave’s latency and amplitude ^[15],^ leading to a reduced slope value.

##### Correlation between AC GDT scores and Temporal Parameters of ABR

There was a positive significant correlation seen between AC GDT scores and the latency of wave I and wave V. These findings indicate that individuals with better AC Gap Detection Thresholds tend to have shorter latencies of wave I and wave V. This shows the importance of neural synchrony in the generation of waves and for GDT.^[12, 16]^

As the latency of wave I is less resistant to changes in repetition rate, the correlation should have been obtained at all repetition rates.^[15]^ The lack of correlation at 90.1/sec could be due to the lower response rate of wave I. In the case of wave V, results should have been similar across rates. However, a positive correlation at higher repetition rates could be due to the higher variability of latency of wave V at these rates and also the large variability in AC GDT values. Thus, it suggests that wave V latency at higher repetition rates is a better predictor of AC GDT.

A negative but non-significant correlation was present between AC GDT and the slope value of wave V. This indicates that individuals with sharper wave V may have better GDT scores. The lack of correlation between the two measures could be due to the different neural mechanisms involved in AC GDT, and in the generation of the slope of wave V. It is believed that neural synchrony is important for AC GDT.^[16]^ Synchronous firing is instrumental in the generation of wave V’s peak, and wave V’s slope and trough depend on the refractory periods of the neural elements involved.^[17]^ Wave V is evoked by neural elements in the lateral lemniscus and inferior colliculus in the midbrain.^[18]^ These elements have differential refractory periods giving rise to a distinct process of temporal resolution. This might have led to a poorer correlation with AC GDT scores.

A significant correlation was visualized between AC GDT scores and the latency parameters of ABR, suggesting a relationship between the behavioral and electrophysiological measures of temporal processing. Hence, measurement of the latency of wave I and wave V of ABR would give an estimate of their AC GDT scores in difficult-to-test populations. AC GDT scores can be predicted better with the latency of wave I and wave V than the slope of wave V. We can infer that AC GDT scores and latency of the wave mainly depend on the synchronous firing of the auditory neurons.The slope of wave V is dependent on the offset and refractory periods of the auditory neurons, leading to contradictory findings for the correlation of these two parameters of ABR with AC GDT scores.

## References

1. Phillips DP, Comeau M, Andrus JN. Auditory Temporal Gap Detection in Children with and without Auditory Processing Disorder. J Am Acad Audiol. 2010;21(6):404–8.

2. Kidd GR, Watson CS, Gygi B. Individual differences in auditory abilities. J Acoust Soc Am. 2007;122(1):418–35.

3. Formby C, Gerber MJ, Sherlock LP, Magder LS. Evidence for an across-frequency, between-channel process in asymptotic monaural temporal gap detection. J Acoust Soc Am. 2002;103(6):3554–60.

4. Walker KM, Brown DK, Scarff C, Watson C, Muir P, Phillips DP. Temporal Processing Performance, Reading Performance, and Auditory Processing Disorder in Learning-Impaired Children and Controls. Can J Speech-Language Pathol Audiol. 2011;35(1):6–17.

5. Jacobson J. An overview of the auditory brainstem response [Internet]. College-Hill Press, San Diego. College-Hill Press; 1985 [cited 2019 Mar 29]. 3–12 p. Available from: https://ci.nii.ac.jp/naid/10010276545/

6. Venkatesan S. Ethical guidelines for bio-behavioural research involving human subjects. Dr. Vijayalakshmi Basavaraj, Director, All India Institute of Speech and Hearing, Manasagangothri, Mysore; 2009.

7. ANSI. ANSI/ASA S3.1-1999 - Maximum Permissible Ambient Noise Levels for Audiometric Test Rooms [Internet]. New York: American National Standards Institute. 2013 [cited 2019 Mar 29]. Available from: https://webstore.ansi.org/standards/asa/ansiasas31999r2013

8. Kwon BJ. AUX: A scripting language for auditory signal processing and software packages for psychoacoustic experiments and education. Behav Res Methods [Internet]. 2012 Jun 20 [cited 2019 Mar 29];44(2):361–73. Available from: http://www.springerlink.com/index/10.3758/s13428-011-0161-1

9. Levitt H. Transformed Up-Down Methods in Psychoacoustics. J Acoust Soc Am [Internet]. 1971 Feb 3 [cited 2019 Mar 29];49(2B):467–77. Available from: http://asa.scitation.org/doi/10.1121/1.1912375

10. Gopal K V., Kowalski J. Slope analysis of auditory brainstem responses in children at risk of central auditory processing disorders. Scand Audiol. 1999;28(2):85–90.

11. Hess BA, Blumsack JT, Ross ME, Brock RE. Performance at different stimulus intensities with the within-and across-channel adaptive tests of temporal resolution. Int J Audiol. 2012;51(12):900–5.

12. Rowe MJ. Normal variability of the brain-stem auditory evoked response in young and old adult subjects. Electroencephalogr Clin Neurophysiol. 1978;44(4):459–70.

13. Stockard JE, Stockard JJ, Westmoreland BF, Corfits JL. Brainstem Auditory-Evoked Responses. Arch Neurol [Internet]. 1979;36(13):823. Available from: http://archneur.jamanetwork.com/article.aspx?doi=10.1001/archneur.1979.00500490037006

14. Sumner CJ, Palmer AR. Auditory nerve fibre responses in the ferret. Eur J Neurosci [Internet]. 2012 Aug 1 [cited 2019 Mar 12];36(4):2428–39. Available from: http://doi.wiley.com/10.1111/j.1460-9568.2012.08151.x

15. Fowler CG, Noffsinger D. Effects of Stimulus Repetition Rate and Frequency on the Auditory Brainstem Response in Normal, Cochlear-Impaired, and VIII Nerve/Brainstem-Impaired Subjects. J Speech, Lang Hear Res [Internet]. 1983 Dec [cited 2019 Mar 13];26(4):560–7. Available from: http://pubs.asha.org/doi/10.1044/jshr.2604.560

16. Zeng FG, Oba S, Garde S, Sininger Y, Starr A. Temporal and speech processing deficits in auditory neuropathy. Neuroreport [Internet]. 1999 Nov 8 [cited 2019 Mar 12];10(16):3429–35. Available from: http://www.ncbi.nlm.nih.gov/pubmed/10599857

17. Don M, Allen AR, Starr A. Effect of Click Rate on the Latency of Auditory Brain Stem Responses in Humans. Ann Otol Rhinol Laryngol [Internet]. 1977 Mar 29 [cited 2019 Mar 12];86(2):186–95. Available from: http://journals.sagepub.com/doi/10.1177/000348947708600209

18. Jewett DL, Romano MN, Williston JS. Human auditory evoked potentials: possible brain stem components detected on the scalp. Science [Internet]. 1970 Mar 13 [cited 2019 Mar 13];167(3924):1517–8. Available from: http://www.ncbi.nlm.nih.gov/pubmed/5415287

